# Remote organ cancer adversely alters renal function and induces kidney injury, inflammation, and fibrosis

**DOI:** 10.1101/2024.07.29.605635

**Authors:** Dana Hammouri, Andrew Orwick, Mark Doll, Dianet Sanchez Vega, Parag P. Shah, Christopher J. Clarke, Brian Clem, Levi J. Beverly, Leah J. Siskind

## Abstract

Approximately 30% of cancer patients experience kidney complications, which hinder optimal cancer management, imposing a burden on patients’ quality of life and the healthcare system. The etiology of kidney complications in cancer patients is often attributed to nephrotoxic oncological therapies. However, the direct impact of cancer on kidney health is underestimated, as most nephrotoxic oncological therapies have been studied in animal models that do not have cancer. Our previous study demonstrated that metastatic lung cancer adversely alters kidney physiology and function, and exacerbates chemotherapy-induced nephrotoxicity, indicating lung cancer-kidney crosstalk. The current study examines whether this phenomenon is specific to the employed cancer model. Female and male mice of various strains were injected with different cell lines representing human and mouse lung cancer, breast cancer, and melanoma, and their kidney tissues were analyzed for toxicity and fibrosis. The impact of cancer on the kidney varied by cancer type. Breast cancer and specific subtypes of lung cancer, including KRAS- and EGFR-mutant cancer, pathologically altered kidney physiology and function in a manner dependent on the metastatic potential of the cell line. This was independent of mouse strain, sex, and cancer cell line origin. Moreover, tumor DNA was not detected in the renal tissue, excluding metastases to the kidney as a causative factor for the observed pathological alterations. Lewis lung carcinoma and B16 melanoma did not cause nephrotoxicity, regardless of the tumor size. Our results confirm cancer-kidney crosstalk in specific cancer types and highlight gaps in understanding the risk of renal complications in cancer patients. In the era of precision medicine, further research is essential to identify at-risk oncology populations, enabling early detection and management of renal complications.

## Introduction

The increasing prevalence of kidney disease among cancer patients has given rise to the field of onconephrology, an emerging subspecialty dedicated to addressing the complex intersections between cancer and kidney disease [1]. Approximately 30% of cancer patients experience kidney complications, with 8% to 34% of critically ill patients requiring renal replacement therapy [2, 3]. The spectrum of renal disease experienced by cancer patients is broad, encompassing acute kidney injury (AKI), chronic kidney disease (CKD), proteinuria, nephrotic syndrome, nephritis, and complex electrolyte disorders [4-9]. The concomitant manifestations of renal complications in cancer patients are frequently attributed to the nephrotoxic effects of antineoplastic therapies, potentially compromising the eligibility for optimal cancer treatments and significantly imposing a burden on patients’ quality of life [6, 10, 11].

Chemotherapy remains a cornerstone in the treatment of many types of cancer and has markedly enhanced remission rates. However, its association with nephrotoxicity is extensively documented, with cisplatin (CDDP) recognized as one of the most nephrotoxic agents used to treat several malignancies [12, 13]. Thirty percent of cancer patients treated with CDDP develop AKI, a rapid decline in kidney function. AKI not only contributes to substantial morbidity but also increases the risk of chronic kidney disease (CKD), a leading cause of mortality worldwide [14-18]. The etiology of kidney complications in cancer patients receiving CDDP is poorly understood and is often attributed solely to the administration of CDDP. This oversimplified view likely stems from the reliance of preclinical studies of CDDP-induced nephrotoxicity (CIN) on non-cancer animal models, neglecting the potential role of cancer contributing to nephrotoxicity. This may partly account for the translational failure observed with candidate drugs that, while effective in preclinical non-cancer models of CIN, have not succeeded clinically. This discrepancy highlights a critical gap in our understanding and indicates that factors beyond CDDP are not adequately considered in the preclinical models. Moreover, it raises the question of whether the high incidence of AKI observed in cancer patients treated with CDDP is solely due to the nephrotoxic mechanism of the drug or whether it’s influenced by the underlying malignancy, which may directly or indirectly contribute to tissue damage, exacerbating CIN. Therefore, integrative research models encompassing cancer pathology and antineoplastic-induced nephrotoxicity are needed. Such models would provide a comprehensive understanding of the multifaceted interactions at play, enhancing the clinical applicability and improving the predictive efficacy of potential nephroprotective candidates in clinical settings.

To address this, we have developed a more clinically relevant model in which tumor-bearing mice receive weekly administration of low-dose CDDP over a four-week period. This model features a syngeneic subcutaneous xenograft of metastatic Kras-mutant, p53 null non-small cell lung carcinoma (NSCLC) in B6;129 F1 hybrid mice [19], potentially representing the most common oncogenic mutations in NSCLC patients receiving CDDP with the highest incidence of AKI compared with other solid organ malignancies [20-24]. Our previous study conclusively demonstrated that metastatic lung cancer significantly disrupts renal physiology, compromises renal function, and exacerbates CDDP-induced nephrotoxicity [19]. Our data support the existence of lung cancer-kidney crosstalk and provide a rationale for further investigation into whether this phenomenon is unique to the lung cancer model employed in our previous study.

In the current study, we expanded the assessment of cancer-kidney crosstalk to encompass other organ malignancies, including diverse variants and stages. Our findings demonstrate a pathological impact of certain types of cancer on the kidney. Triple-negative breast cancer (TNBC) and specific subtypes of lung cancer, including KRAS- and EGFR-mutant cancers, induced kidney injury, inflammation, and fibrosis. Conversely, other cancer types such as Lewis lung carcinoma, and B16 melanoma, were not associated with clear signs of nephrotoxicity.

Our findings support the existence of cancer-kidney crosstalk while highlighting substantial gaps in the current understanding of renal complications in cancer patients. Furthermore, this study emphasizes the urgent need to identify reliable biomarkers to enable early detection of renal complications and ultimately mitigate severe clinical outcomes.

## Methods

### Animals and animal care

All mice were housed under standard laboratory conditions, with a 12-hour light/dark cycle provided with food and water ad libitum. All animal experimental procedures were approved by the Institutional Animal Care and Use Committee of the University of Louisville or the Institutional Animal Care and Use Committee of Stony Brook University, adhering to the American Veterinary Medical Association’s guidelines.

### Tumor Outgrowth Studies

#### Syngeneic xenograft model of Kras^G12D^ Trp53^KO^ lung cancer

Eight-to ten-week-old male B6129SF1/J (B6;129) mice were purchased from the Jackson Laboratory (Strain #:101043). Three previously developed mouse cell lines of Kras^G12D^ Trp53^KO^ NSCLC were used [25]. This includes two cell lines derived from lymph node metastases (238N1 and 482N1) and one from a primary lung tumor (394T4). Twenty mice were randomly assigned into four groups: non-cancer and three cancer groups. Mice in the cancer groups received a subcutaneous injection of 1 × 10^4^ cancer cells into the right rear flank, as previously described [19]. As for the IV-Kras-mutant lung cancer model, 1 × 10^4^ of 238N1 or 394T4 were injected in the tail vein of eight-to ten-week-old male B6129 mice. Tumor progression was monitored, and mice were euthanized 28 days post-injection or when mice reached the endpoint as per the IACUC protocol and as specified in the manuscript.

#### Immunocompromised xenograft model of human KRAS^G12C^ Lung cancer (A549)

NOD.Cg-Rag1tm1Mom Il2rgtm1Wjl Tg (CMV-IL3, CSF2, KITLG)1Eav/J (NRGS) mice were purchased from the Jackson Laboratory (Strain #: 024099). A549 human NSCLC cell line was purchased from ATCC (Rockville, MD, USA) and verified via short tandem repeat (STR). 1×10^6^ cells were subcutaneously implanted in 6-to 10-week-old male and female mice. Mice were monitored and euthanized when tumors reached designated endpoints, usually around 40 days post-cancer cell injection.

#### Immunocompromised xenograft model of human EGFR-mutant Lung cancer (PC9)

NOD.Cg-Prkdcscid Il2rgtm1Wjl/SzJ (NSG) mice were purchased from the Jackson Laboratory (Strain #: 005557). PC9 human NSCLC cells were purchased from ATCC and verified via STR. Cells were maintained according to the manufacturer’s recommendations. 1×10^6^ cells were subcutaneously implanted in 6-to 10-week-old male and female mice. Mice were monitored and euthanized when tumors reached designated endpoints, usually around 40 days post-cancer cell injection.

#### Immunocompromised xenograft model of human breast cancer

Female NSG mice at 8 weeks age were purchased from Jackson Laboratories. MDA-MB-231 human breast cancer cells were purchased from ATCC (Rockville, MD, USA), cultured in DMEM medium supplemented with 10% fetal bovine serum (Invitrogen, Carlsbad, CA, USA), and tested monthly for mycoplasma contamination. For orthotopic experiments, 2 × 10^6^ cells in 25ul PBS were mixed in a ratio of 1:1 with Matrigel and orthotopically implanted in the 4^th^ mammary fat pad. Mice were regularly monitored for tumor development and once palpable tumors had formed (typically 7-10d), primary tumor size was monitored with caliper measurements twice a week. When tumors reached designated endpoints (typically 35-40d post injection), animals were euthanized for tissue collection.

#### Syngeneic xenograft model of mouse breast cancer

Female Balb/c mice at 8 weeks of age were purchased from Jackson Laboratories. 4T1 mouse breast cancer cells were purchased from ATCC (Rockville, MD, USA), maintained in RPMI medium supplemented with 10% FBS, and routinely subcultured every 3-4 days. For orthotopic experiments, 2.5 × 10^4^ cells in 25ul PBS were mixed in a ratio of 1:1 with Matrigel and orthotopically implanted in the 4^th^ mammary fat pad. Mice were regularly monitored for tumor development and once palpable tumors had formed (typically 5-7d), primary tumor size was monitored with caliper measurements twice a week. When tumors reached designated endpoints (typically 21d post injection), animals were euthanized for tissue collection.

#### Congenic xenograft model of Lewis Lung Cancer (LLC)

Eight-to ten-week-old male B6129SF1/J (B6;129) mice were purchased from the Jackson Laboratory (Strain #:101043). LLC cells were obtained from ATCC (Rockville, MD, USA) and maintained in culture as recommended. Mice received a subcutaneous injection of 1 × 10^6^ LLC cells into the right rear flank, as previously described [19]. Tumor progression was monitored, and mice were euthanized 19 days post-injection when mice reached the endpoint as per the IACUC protocol.

#### Congenic xenograft model of B16 melanoma

Eight-to ten-week-old male B6129SF1/J (B6;129) mice were purchased from the Jackson Laboratory (Strain #:101043). B16 cells were obtained from ATCC (Rockville, MD, USA) and maintained in culture as recommended. Eight-to ten-week-old mice received a subcutaneous injection of 1 × 10^6^ B16 cells into the right rear flank as previously described [19]. Tumor progression was monitored, and mice were euthanized 19 days post-injection when mice reached the endpoint as per the IACUC protocol.

#### Blood Urea Nitrogen and Neutrophil Gelatinase-Associated Lipocalin Determination

Blood urea nitrogen (BUN) was measured in the plasma of mice using AMS Diagnostics detection kit (No. 80146, AMS Diagnostics), per the manufacturer’s instructions. For neutrophil gelatinase-associated lipocalin, ELISA (NGAL; DY1857, R&D Systems) was performed on mouse urine per the manufacturer’s protocol and as previously described [26].

#### Tissue Histology and Immunohistochemistry (IHC)

Renal tissue samples were fixed in 10% neutral formalin and paraffin-embedded (FFPE) for histology and immunohistochemistry (IHC). For quantitative assessment of tubulointerstitial fibrosis, kidney sections (5μm) were stained with Sirius Red/Fast Green (SRFG) (0.1% Sirius red (Sigma, 365548) and 0.1% fast green (Sigma, F7258) in saturated picric acid or PicroSirius red (0.1% Sirius red in saturated picric acid) for 1 h, followed by two washes with 0.5% Glacial acetic acid. Tissue samples were then dehydrated and fixed using histomount mounting solution. The Sirius red–positive area in the kidney section was measured using QuPath 0.5.1.

For Hematoxylin and eosin (H&E) staining, lungs were harvested from mice and were formalin-fixed and paraffin-embedded (FFPE). Lung tissue sections (5μm) were deparaffinized, rehydrated, and then stained with H&E. After staining and dehydration, sections were mounted and scanned for microscopic observation.

For IHC, FFPE sections (5μm) were deparaffinized and rehydrated. Antigen retrieval was performed using citrate buffer (pH 6.0) for KIM-1 staining or Tris-EDTA (PH 9) for Kras^G12D^ staining for 30 min, then cooled to room temperature for 30 minutes. Endogenous peroxidase was blocked by incubation in 3% H_2_O_2_ solution for 30 minutes. This was followed by further blocking using avidin/biotin blocking solution for 20 minutes. Subsequently, the sections were blocked with 10% normal goat or rabbit serum in 1% PBS for 30 minutes. Slides were then incubated with primary antibody diluted in 1% PBS overnight at 4°C. The primary antibody was washed off with 1% PBS three times and the secondary antibody was added for an incubation time of 1 hr. Slides were rinsed with 1% PBS for 5 min, and ABC reagents (PK-4001, Vectastain) were added for 30 min, followed by three washes with 1% PBS. ImmPACT NovaRed peroxidase substrate (Vector SK-4805) was added to slides at room temperature for 2-3 min. Slides were counterstained and mounted as previously described [26]. The following antibodies were used: Anti-Kras^G12D^ (Invitrogen, 536256) and Anti-KIM-1 (R&D Systems, AF1817).

### DNA Extraction

DNA was extracted from lung and kidney tissue with the E.Z.N.A Tissue DNA Kit (OMEGA) according to the manufacturer’s protocol. DNA was quantified using NanoDrop 2000c spectrophotometer (Thermo Scientific, ND2000CLAPTOP).

#### Lung Metastases Detection by Droplet Digital PCR (ddPCR)

ddPCR was performed using the QX200 Droplet Digital PCR system (Bio-Rad Laboratories). Samples were prepared by mixing 12.5 µL ddPCR Supermix for probes (No dUTP, Bio-Rad Laboratories), 1.25 µL ddPCR™ probe assay Kit (Bio-Rad Laboratories) which consist of forward (CGCAATCCTTTATTCTGTTCGA) and reverse (GAGACGGGTCTTGCTATTGTAGCTA) PCR primers and FAM-labeled fluorescent probe specific for the mutation assay (ATCGATAAGCTTGATATCGAATT), and 11 µL of template DNA in a final reaction volume of 25 μL. Droplets were generated by a QX200 droplet generator. Endpoint PCR was performed on a C1000 Touch Thermal Cycler (Bio-Rad Laboratories). Thermal cycling profile for Kras^G12D^ mutation assay started with a hot start denaturation step of 10 mins at 95°C, followed by 40 cycles of 94°C for 30 s, 60°C for 1 min. These cycles were followed by 98°C for 10 mins and then 12°C hold. Then, PCR products were loaded into the QX200 droplet reader and analyzed by QuantaSoft version 1.7.4.0917 (Bio-Rad Laboratories). Data were reported as copies per 1 μg. For each assay, water without templates served as a control for detecting environmental contamination; a negative control (genomic DNA from a non-cancer mouse) was used to estimate the false-positive rates; and a positive control containing genomic DNA from kras mutant mouse was used to verify the assay performance and determine the threshold value of fluorescent signals.

#### Protein Quantification and Western Blot Analysis

Kidney tissues were homogenized in cell extraction buffer (ThermoFisher Scientific) containing a complete protease and phosphatase inhibitor cocktail (ThermoFisher Scientific, A32959). Homogenates were centrifuged at 15,000 g for 10 min at 4 C. Supernatants were removed and protein concentrations were determined using BCA reagent (ThermoFisher Scientific). Forty micrograms of kidney homogenate protein were denatured at 95°C for 5 min, after which they were loaded and separated on 4–12% gradient Tris-glycine-SDS polyacrylamide gels. Protein was then transferred to PVDF membrane, which was then incubated in 5% (w/v) dried milk in Tris-buffered saline-0.1% Tween 20 (TBST) for 1 h. Membranes were incubated with primary antibody (1:1000 in 1% dried milk) overnight at 4°C, followed by three times wash with TBST. Membranes were then incubated with secondary antibodies conjugated with horseradish peroxidase (1:20000) in TBST containing 1% (w/v) dried milk. Following two washes with 1% (wt/vol) dried milk and one wash in TBST, membrane proteins were detected by chemiluminescence substrate. The following antibodies were used: Anti-fibronectin (Millipore, AB1954), TGFb (Abcam, ab215715), α-SMA (Abcam, ab5694), B-Catenin (Cell signaling, 8480), Vimentin (Cell signaling,5741), Claudin-1 (Cell signaling, 13255), Tenascin C (Abcam, ab108930), Snail (Cell signaling, 3879), α-tubulin (Santa Cruz Biotechnology, sc-5286,).

### Gene Expression

RNA was isolated from kidney tissue using E.Z.N.A. Total RNA Kit 1 (OMEGA) per the manufacturer’s protocol. cDNA was synthesized with High-Capacity cDNA. Reverse Transcriptase PCR (ThermoFisher Scientific) per the manufacturer’s instructions. Gene-specific cDNA was quantified with real-time quantitative PCR using predesigned TaqMan or self-designed SYBR assays. The following TaqMan primers were purchased from ThermoFisher Scientific: tumor necrosis factor-a (Tnf-α; Mm00443258_m1), chemokine (C-X-C motif) ligand 1 (Cxcl1; Mm04207460_m1), IL-6 (Mm00446190_m1), monocyte chemoattractant protein-1 (MCP-1; Mm00441242_m1), and the housekeeping gene b2-microglobulin (B2m; Mm00437762_m1). The following primers shown were self-designed: kidney injury molecule-1 (Kim-1) forward: AGATCCACACATGTACCAACATCAA and reverse: CAGTGCCATTCCAGTCTGGTTT, tissue inhibitor of metalloproteinase (Timp-1) forward: GCAACTCGGACCTGGTCATAA and reverse: TTAGTCATCTTGATCTTATAACGCTGGTA, NLR family pyrin domain-containing 3 (Nlrp3) forward: AAGATGAAGGACCCACAGTGTAACTT and reverse: CAGATTGAAGTAAGGCCGGAATT, collagen type I-a1 (Col1a1) forward: CGATGGATTCCCGTTCGAGTA and reverse: GTGGACATTAGGCGCAGGAA, Low density lipoprotein receptor-related protein 3 (Lrp-3) forward: AAAATGGAAACGGGGTGACTT and reverse: GGCTGCATACATTGGGTTTTC3 and Klotho forward: GTACCTGGTTGCCCACAACCTA and reverse: GCGGAAAGAGGTGTTGTAGAGATG. Quantitative RT-PCR was done with either iTaq Universal Probes Supermix (No. 172–5134, Bio-Rad) or iTaq Universal SYBR Green Supermix (No. 172–5124, Bio-Rad).

### Statistical analysis

Data are expressed as mean ± SD or median ± range for all experiments. Homogeneity of variance was assessed with F-Test or Bartlett’s Test. Single comparisons of normally distributed continuous data were analyzed using a one-sided unpaired t-test as appropriate. Multiple comparisons of normally distributed data were analyzed by one-way ANOVA, and group means were compared using Tukey post-hoc when the variance was homogeneous; otherwise, Kruskal-Wallis test was used, followed by Dunn’s multiple comparison test. The criterion for statistical differences was *p<0.05, **p<0.01, ***p<0.001, ****p<0.0001.

## Results

### Kras^G12D^ Trp53^KO^ lung cancer induces kidney injury and fibrosis and adversely affects renal function in a manner dependent on its metastatic potential

Our previous study demonstrated that mice subcutaneously injected with the Kras^G12D^, Trp53^KO^-metastatic lung cancer cell line 238N1 exhibited altered renal physiology, impaired kidney function, and increased levels of CIN nephrotoxicity [27]. This phenomenon prompted us to investigate whether this pathological crosstalk is exclusive to the lung cancer type that the 238N1 cell line represents. Therefore, we evaluated kidney physiology and function in the presence of different cell lines of Kras^G12D^ Trp53^KO^ lung cancer with varying metastatic potential [25]. This includes a highly metastatic cell line (238N1), a less aggressive metastatic cell line (482N1), and a cell line with a low metastatic potential (394T4). Cells were subcutaneously injected into the right rear flank of mice. Mice were followed for tumor size and were euthanized 28 days following cancer cell injection or once the tumor size reached the endpoint per IACUC protocol **(Fig. 1A, 1B)**. Mice injected with the 238N1 and 438N1 cell lines exhibited comparable average tumor weight and volume, whereas the 394T4 group demonstrated smaller tumor size, suggesting a slower growth rate and/ or less aggressive nature of this cell line **(Fig. 1C-1E)**. Using histological staining, we confirmed the local metastatic potential of the cell lines in lung tissue. The 238N1 group exhibited a higher level of lung metastases compared to the 482N1 and 394T4 groups **(Fig. 1F)**. This was associated with an increased level of blood urea nitrogen (BUN) and urinary neutrophil gelatinase-associated lipocalin (NGAL), indicating kidney function deterioration and tissue damage, respectively **(Fig. 2A, 2B)**. Although to a lesser extent, we observed an increase in the level of urinary NGAL in the 482N1 and 394T4 groups, suggesting a mild degree of kidney injury. This 238N1 group had a significantly higher average increase in tubular expression of kidney injury molecule-1 (Kim-1), indicating a pronounced tubular injury compared to the other groups **(Fig. 2C, 2I)**. Quantitative real-time (qPCR) analysis showed a significant decrease in the gene expression of the low-density lipoprotein-related protein 2 (Lrp2) in the cancer groups, implicating an injury to the tubular basement membrane and loss of the brush border in the proximal tubules **(Fig. 2D)**. Additionally, we observed an increase in the gene expression of the inflammatory markers in all the cancer groups indicated by the significant increase in the level of the gene expression of certain cytokines and chemokines, such as interleukin 6 (IL-6), chemokine (C-X-C motif) ligand 1 (Cxcl1), monocyte chemoattractant protein-1 (Mcp-1), and tumor necrosis factor (Tnf-α) **(Fig. 2E-2H)**. Notably, the levels of Mcp-1 and Tnf-α were significantly higher in the 394T4 group than in the other groups, potentially indicating a delayed injury/inflammatory response.

**Figure 1.**
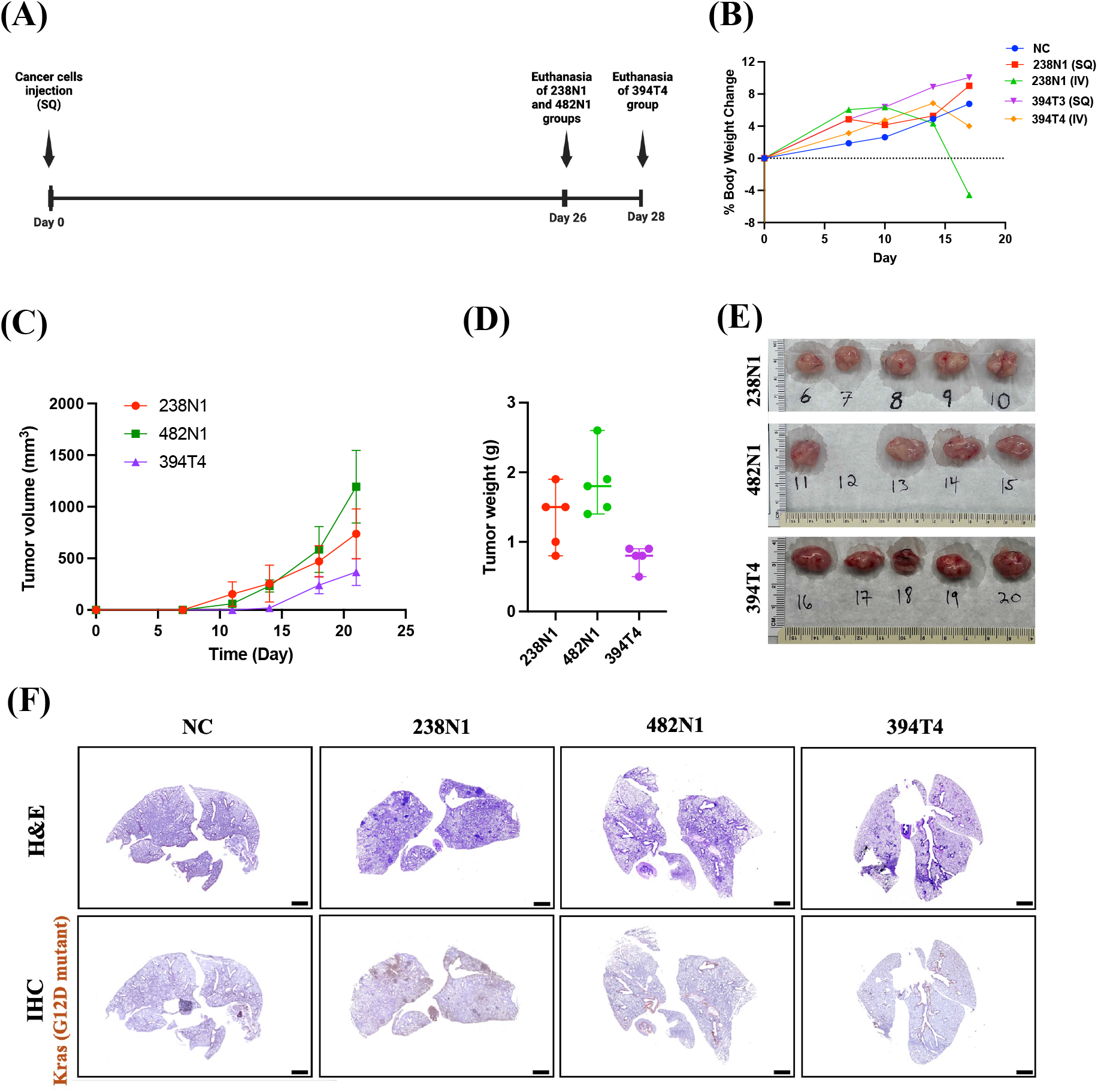
Validation of the subcutaneous implantation in the syngeneic xenograft models of Kras^G12D^ Trp53^KO^ lung cancer: **(A)** Experimental design of lung cancer models in B6;129 male mice subcutaneously, injected with 1 × 10^4^ of different cell lines of Kras^G12D^ Trp53^KO^ NSCLC. **(B)** Percentage change in body weight. **(C)** Tumor volume and **(D)** weight. **(E)** Images of tumors following euthanasia. The image of tumor sample 12 could not be obtained due to the loss of the sample prior to imaging. **(F)** Representative images of Hematoxylin and eosin (H&E) and immunohistochemistry (IHC) staining of Kras^G12D^ mutant in the lungs of mice with or without cancer. Scale size: 1000 µm. Data are presented as median ± range. n = 5. Non-Cancer (NC).

**Figure 2.**
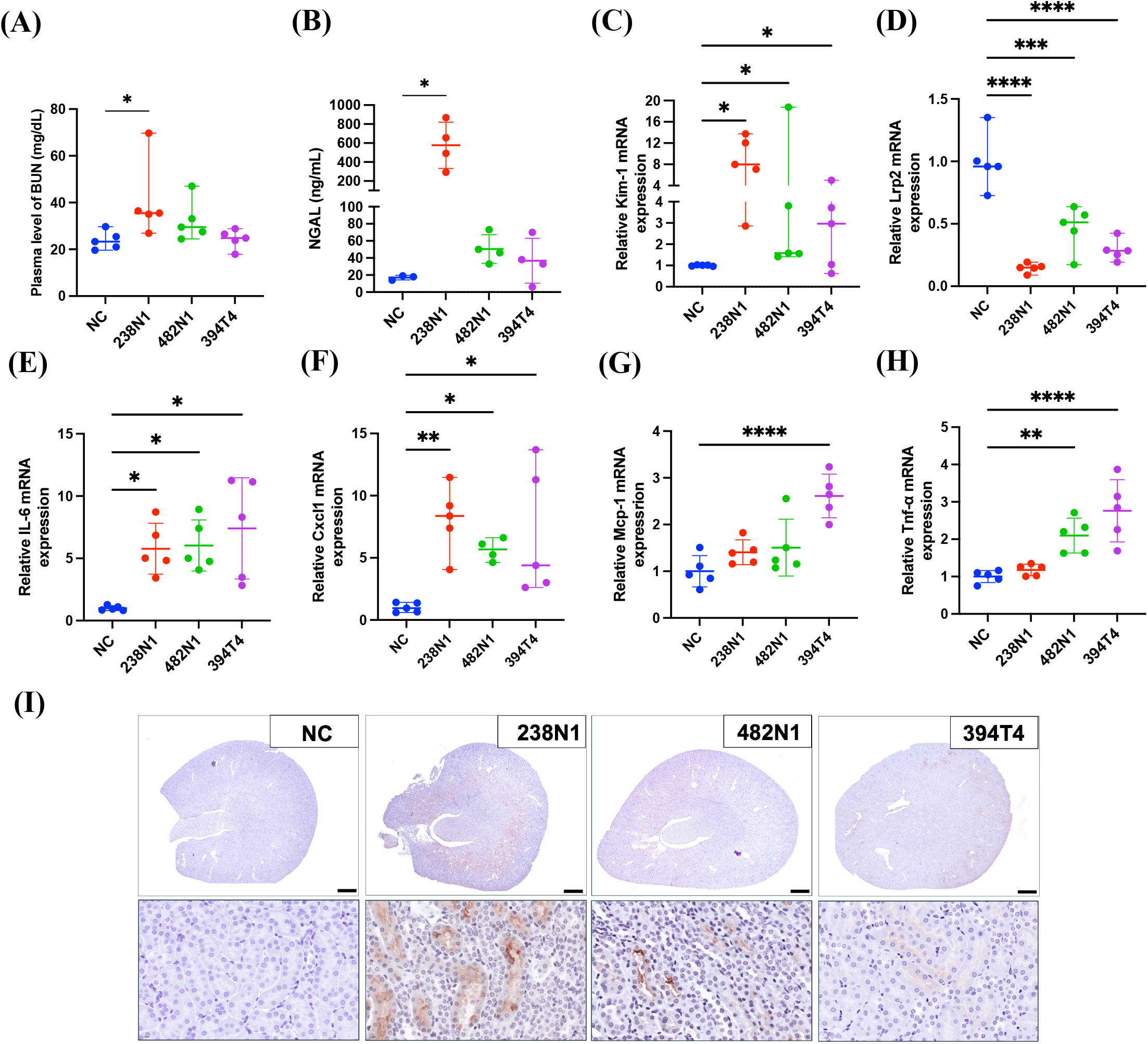
Kras^G12D^ Trp53^KO^ lung cancer adversely affects renal function, induces kidney injury and triggers an inflammatory response in a manner dependent on its metastatic potential. **(A)** Scatter plot of blood urea nitrogen (BUN) **(B)** urinary NGAL. mRNA level of **(C)** Kim-1, **(D)** Lrp2, **(E)** IL-6, **(F)** Cxcl1, **(G)** Mcp-1, **(H)** and Tnf-α. **(I)** Representative images of immunohistochemistry (IHC) staining of KIM-1 protein in kidney sections of B6,129 mice with or without cancer, Scale size: 1000 µm and 50 µm. Data are presented as mean ± SD or median ± range. *P < 0.05, **p<0.01, ***p<0.001 ****p<0.0001, based on one way ANOVA or Kruskal-Wallis test.

To further investigate the impact of Kras^G12D^ Trp53^KO^-mutant lung cancer on renal physiology, we assessed the fibrosis level in kidney tissue. We found a decrease in the gene expression of the renal fibrosis inhibitory factor, Klotho, across all cancer groups, with the most pronounced reduction observed in the 238N1 group [28] **(Fig. 3A)**. Moreover, the 238N1 group had elevated expression of the pro-fibrotic genes, tissue inhibitor of metalloproteinase 1 (Timp-1) and collagen type 1 (Col1a1) **(Fig. 3B, 3C)**. Additionally, the 238N1group showed the highest increase in the levels of the extracellular matrix protein fibronectin and the fibrotic cytokine transforming growth factor beta (TGF-β) in kidney tissue **(Fig. 3E)**. To further determine the extent of fibrosis, we assessed the total collagen content in kidney tissue using picrosirius red staining. The levels of tubulointerstitial collagen were significantly increased in the kidney of the 238N1 and 482N1 groups, with no significant change observed in the 394T4 group **(Fig. 3D, 3F)**. These results collectively demonstrate that Kras^G12D^ Trp53^KO^-mutant lung cancer induces pathological renal alterations dependent on the cancer’s metastatic potential.

**Figure 3.**
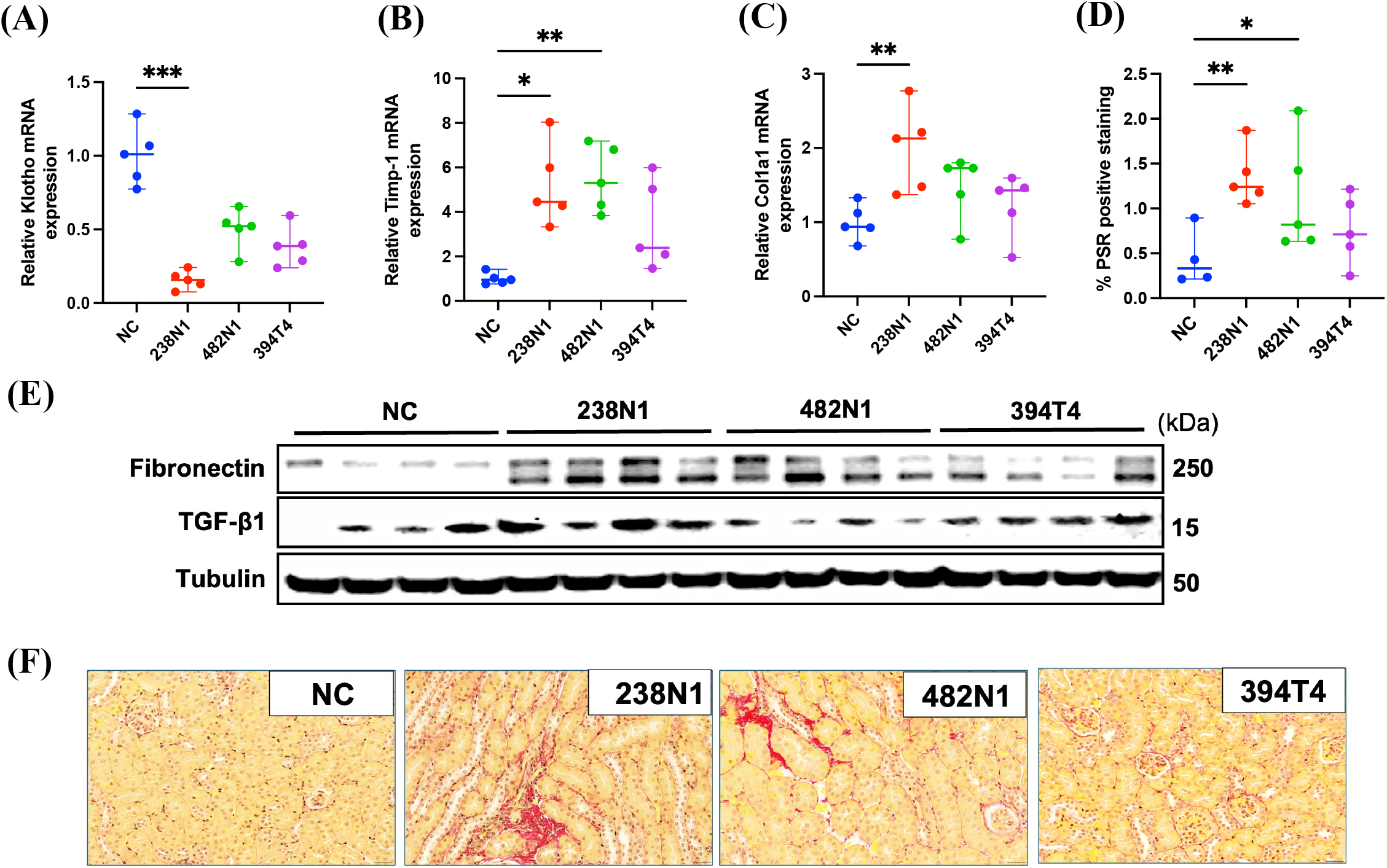
Kras^G12D^ Trp53^KO^ lung cancer induces kidney fibrosis in a stage-dependent manner. Scatter plots of relative mRNA level of **(A)** Klotho, **(B)** Timp-1 and **(C)** Col1a1 in mice with or without cancer. **(D)** Quantification of total picrosirius red (PSR) positive staining of the kidney sections. **(E)** Western blot analysis. **(F)** Representative images of PSR staining. Scale size: 20 µm. Data are presented as mean ± SD or median ± range. *P < 0.05, **p<0.01, ***p<0.001, based on one way ANOVA or Kruskal-Wallis’s test.

### Kras^G12D^ Trp53^KO^ lung cancer is sufficient to induce kidney injury independent of lung injury

Our data indicate that the nephrotoxicity induced by Kras^G12D^ Trp53^KO^ lung cancer is contingent on the metastatic potential of the lung cancer line, implying that local pulmonary tissue damage may be the primary driver of the perceived renal pathology. This observation aligns with the growing body of evidence supporting a bidirectional causality between acute lung injury (ALI) and acute kidney injury (AKI), thereby complicating the interpretation of our results [29, 30]. This raises critical questions regarding whether the pathological renal alterations in mice with Kras^G12D^ Trp53^KO^ lung cancer may be attributable to subsequent lung injuries following lung metastasis. To address this concern, we intravenously administered cancer cells of the highly metastatic cell line (238N1-IV) and the low metastatic cell line (394T4-IV) into the tail vein of mice. This approach ensures the injected cancer cells encounter and seed the lungs [31, 32]. We assessed kidney function and structure, comparing these groups to non-cancer mice and mice in parallel injected subcutaneously with these same cancer cell lines (238N1-SQ and 394T4-SQ). Mice were monitored and euthanized 17 days post-injection, as the 238N1-IV group reached the experimental endpoints per IACUC protocol **(Fig. 4A, 4B)**. The 238N1-SQ group had a larger tumor size than the 394T4-SQ group at 17 days, although the difference was less pronounced than previously observed at 28 days post-injection **(Fig. 4C-4E)**. To validate lung metastases, we analyzed lung tissue using droplet digital PCR (ddPCR). The results demonstrate a significant elevation in tumor DNA levels in the lung tissues of both the 238N1-IV and 394T4-IV groups, thereby confirming the successful seeding of cancer cells in the lungs of mice **(Fig. 4F)**. The 238N1-SQ and 394T4-SQ groups showed comparably low levels of lung metastases **(Fig. 4F)**. Histological staining supported these findings, revealing a significant increase in lung metastases in the 238N1-IV and 394T4-IV groups compared to the other groups **(Fig. 4G)**. Upon evaluating kidney function, we did not observe any significant changes in the level of BUN **(Fig. 5A)**. This could be attributed to the shorter experimental timeframe of 17 days, as opposed to the 28-day period in our previous study, where we noted a significant reduction in kidney function. Conversely, there was an increase in the level of urinary NGAL in both groups of mice subcutaneously and intravenously injected with the highly metastatic cancer cells, 238N1-SQ and 238N1-IV, respectively **(Fig. 5B)**. The tubular expression of Kim-1 was significantly elevated in the 238N1-IV and 394T4-IV suggesting that there could be lung injury causing kidney injury **(Fig. 5C, 5I)**. Although not statistically significant, a slight increase in Kim-1 level was also observed in the 238N1-SQ group at 17 days. qPCR analysis showed a decrease in the gene level of Lrp2 in the cancer groups, suggesting tubular basement membrane injury **(Fig. 5D)**. Clear signs of inflammation were evident in the cancer groups, except for the 394T4-SQ group, as indicated by elevated levels of NOD-like receptor protein 3 (Nlrp3), Cxcl1, Mcp-1, and Tnf-α **(Fig. 5E-5H)**. To rule out the possibility that the metastases to the kidney are contributing to the observed renal pathology, we performed ddPCR analysis on renal tissue from cancer-bearing mice. The analysis revealed no detectable tumor DNA in the renal tissue **(Figure 4F)**.

**Figure 4.**
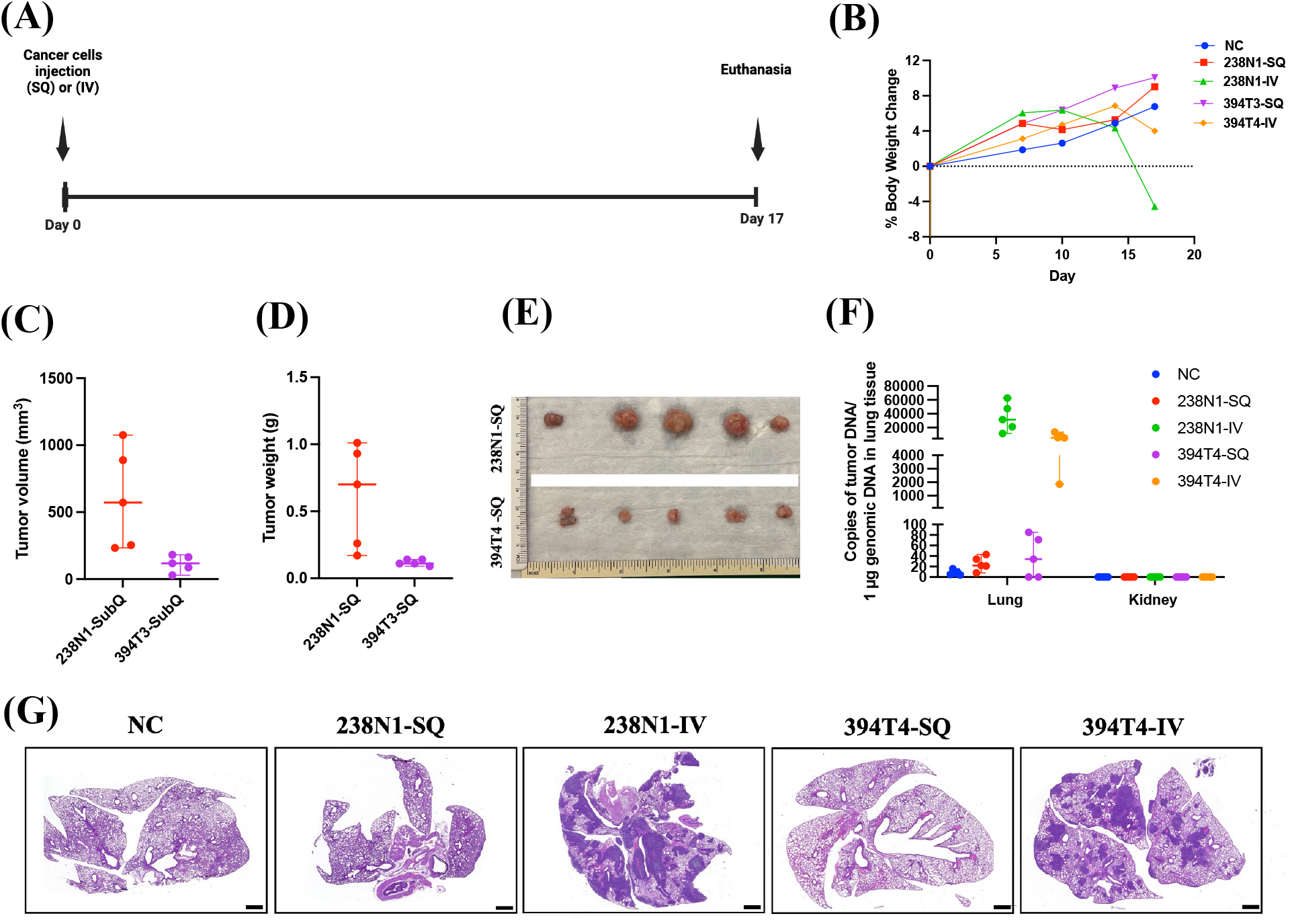
Validation of the subcutaneous and intravenous implantation in the syngeneic xenograft models of Kras^G12D^ Trp53^KO^ lung cancer. **(A)** Experimental design of lung cancer models in B6129 male mice, subcutaneously or intravenously injected with 1 × 10^4^ of 238N1 or 394T4 cell lines. **(B)** Percentage change in body weight. **(C)** Tumor volume and **(D)** weight of mice subcutaneously injected with cancer cells. **(E)** Images of tumors of mice subcutaneously injected with cancer cells following euthanasia. **(F)** Droplet digital PCR (ddPCR) analysis shows the absolute values for DNA copies/1μg of lung or kidney tissue of mice with or without cancer. **(G)** Representative images of H&E staining of lung sections. Scale size: 1000 µm. Data are presented as mean ± SD or median ± range. n=5

**Figure 5.**
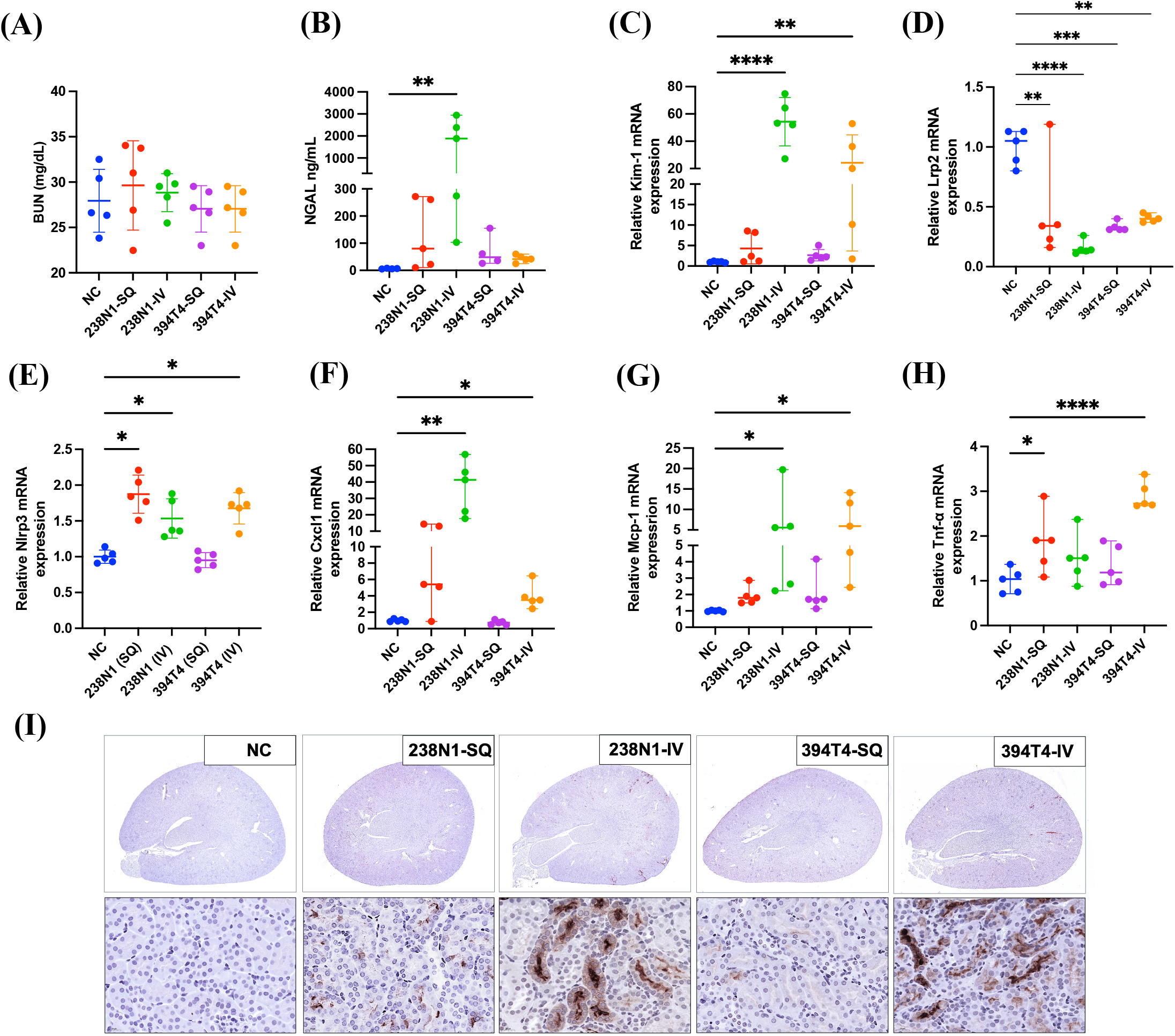
Metastatic Kras^G12D^ Trp53^KO^ lung cancer induces kidney injury and inflammation independent of lung injury. **(A)** Scatter plot shows the level of BUN and **(B)** NGAL. **(C)** mRNA level of Kim-1, **(D)** Lrp2, **(E)** Nlrp3, **(F)** Cxcl1, **(G)** Mcp-1, **(H)** and Tnf-α. **(I)** Representative images of IHC staining of KIM-1 in kidney sections of mice with or without cancer, Scale size: 1000 µm and 50 µm. Data are presented as mean ± SD or median ± range. *P < 0.05, **p<0.01, ***p<0.001 ****p<0.0001, based on one way ANOVA or Kruskal-Wallis test.

Our observations in the intravenously injected mice reinforce the accumulating evidence of the pathological crosstalk between lung and kidney injuries. Moreover, our results demonstrate that the subcutaneous 238N1 tumor alone is sufficient to induce kidney injury and trigger an inflammatory response independent of lung metastases or direct lung injury.

### Kras^G12D^ Trp53^KO^-mutant lung cancer induces tubulointerstitial fibrosis independent of lung injury

To further investigate the renal pathology in mice subcutaneously or intravenously injected with cancer cells, we evaluated the extent of fibrosis in renal tissue. We observed a significant decrease in the gene expression of Klotho in the kidney tissue of both the 238N1-SQ and 238N1-IV groups. Whereas the 394T4-SQ and 394T4-IV groups showed a less pronounced reduction compared to the 238N1 groups **(Fig.6A)**. Additionally, the interstitial collagen content in the kidney tissue was significantly elevated in the 238N1-SQ and 238N1-IV groups, with no significant changes observed in the 394T4-SQ or 394T4-IV groups **(Fig. 6B, 6D)**. Furthermore, the levels of fibronectin and TGF-β were markedly increased in the 238N1-SQ and 238N1-IV groups but not the 394T4-SQ or 394T4-IV groups **(Fig. 6C)**. Given that epithelial-mesenchymal transition (EMT) directly contributes to the fibroblast pool during fibrogenesis, we assessed the protein levels of EMT markers in the kidney tissue [33, 34]. We found a significant upregulation of EMT markers, including β-Catenin, vimentin, and claudin-1, exclusively in the 238N1-SQ and 238N1-IV groups **(Fig. C)**. These findings suggest that the renal fibrosis observed in the 238N1 groups is likely due to cancer-kidney crosstalk rather than solely from metastasis to the lungs and induction of lung injury. This is further supported by the observation in the 394T4-IV group, which had elevated markers of kidney injury and inflammation, yet showed no indications of altered EMT or fibrosis in the kidney.

**Figure 6.**
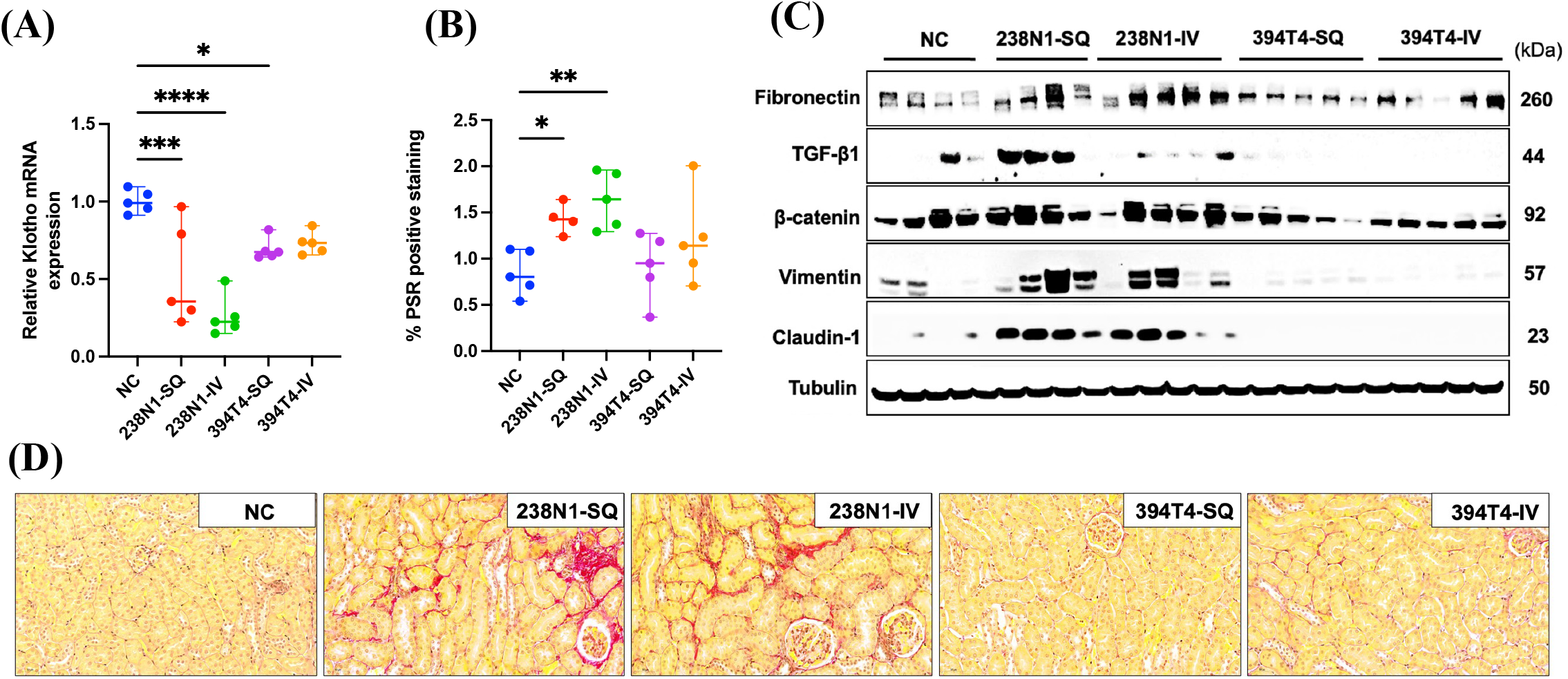
Metastatic Kras^G12D^ Trp53^KO^ lung cancer induces kidney fibrosis independent of lung injury. **(A)** Scatter plot shows the mRNA level of Klotho, **(B)** Quantification of total PSR positive staining of the kidney sections. **(C)** Western blot analysis. **(D)** Representative images of PSR staining. Scale size: 20 µm. **(F)** Western blot analysis. Data are presented as mean ± SD or median ± range. *P < 0.05, **p<0.01, ***p<0.001, based on one way ANOVA or Kruskal-Wallis’s test.

### Human KRAS^G12C^ and EGFR-mutant lung cancer cell lines induce nephrotoxicity

Given our findings that support the existence of cancer-kidney crosstalk associated with Kras^G12D^ Trp53^KO^ NSCLC, we aimed to investigate whether this phenomenon extends to human lung cancer cell lines or other types of lung cancer characterized by different driver mutations and originating from different cell types in various animal strains. To examine whether cancer-kidney crosstalk is specific to mouse Kras^G12D^ -mutant lung cancer in B6;129 mice, we evaluated the renal tissue of NRGS female and male mice bearing human KRAS^G12C^-mutant lung cancer (A549). Cancer mice showed a significant increase in kidney injury, evidenced by elevated tubular expression of KIM-1 **(Fig. 7A)**. Furthermore, these mice showed increased accumulation of interstitial collagen assessed by picrosirius red staining as compared to non-cancer controls, suggesting enhanced renal fibrosis in the presence of cancer. **(Fig. 7A, 7B)**. To examine whether these pathological renal alterations are specific to KRAS mutations, we assessed the kidney tissue of NSG female and male mice bearing a subcutaneous tumor formed by human epidermal growth factor receptor (EGFR)-mutant NSCLC cells (PC9). Consistent with our findings in mice with KRAS^G12C^-mutant lung cancer, the EGFR-mutant lung tumor induced a pathological crosstalk with the kidney, resulting in increased tubular injury as determined by IHC staining of Kim-1 and interstitial fibrosis as determined by picrosirius red staining of collagen **(Fig. 7C, 7D)**. To further evaluate the presence of this crosstalk in different types of lung cancer, we examined kidney injury and fibrosis in B6;129 male mice bearing Lewis lung carcinoma (LLC) tumors. Despite the development of large tumor size **(Fig. S1)**, no significant signs of kidney injury or fibrosis were observed in the renal tissue compared to non-cancer controls **(Fig. 7E, 7F, S1)**. Although urinary NGAL levels were slightly elevated, this increase did not consistently correlate with renal injury or fibrosis assessments **(Fig. S1)**. Given the lack of specificity of NGAL to kidney injury, its elevated level may be attributable to other altered factors in the organ tissues of mice with LLC.

**Figure 7.**
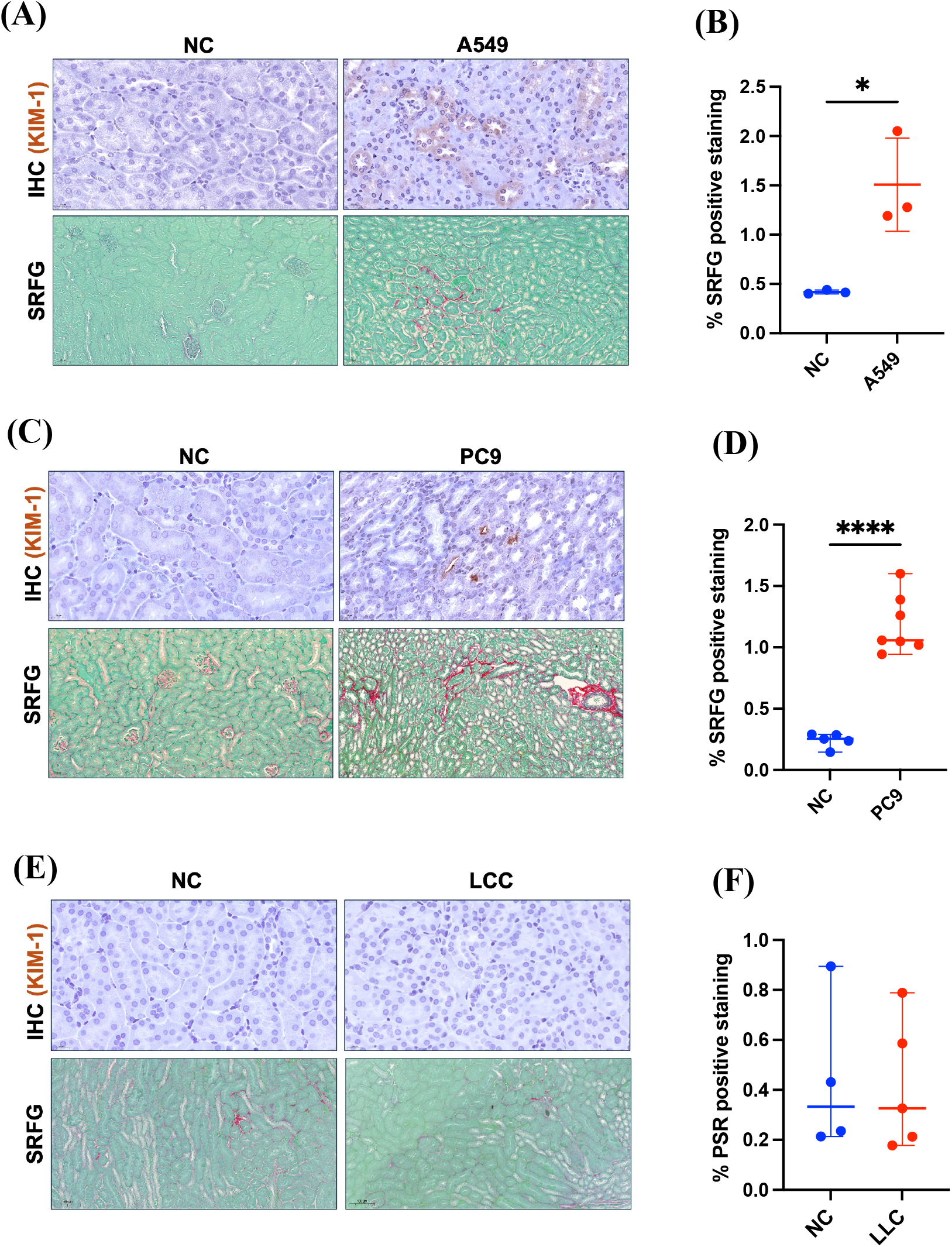
Human KRAS^G12C^-mutant lung cancer (A549) and EGFR-mutant lung cancer (PC9) induce cancer-kidney crosstalk. **(A)** Representative images of IHC staining of KIM-1 and PSR staining of kidney sections of NRGS female and male mice with or without human kras-mutant lung cancer. Scale size: 50 µm and 20 µm. **(B)** Quantification of total PSR positive staining of the kidney sections **(C)** Representative images of IHC staining of KIM-1 and PSR staining of kidney sections of NSG female and male mice with or without human EGFR-mutant lung cancer. Scale size: 50 µm and 20 µm. **(D)** Quantification of total PSR positive staining of the kidney sections. **(E)** Representative images of IHC staining of KIM-1 and PSR staining of kidney sections of B6,129 male mice with or without Lewis lung carcinoma (LLC). Scale size: 50 µm and 20 µm. **(F)** Quantification of total PSR positive staining of the kidney sections Data are presented as mean ± SD. *P < 0.05, ****p<0.0001, based on one way ANOVA test.

### Cancer-kidney crosstalk is not specific to lung cancer

To investigate whether the pathological crosstalk between cancer and the kidney is exclusive to lung cancer, we evaluated kidney tissue in the presence of cancer cell lines derived from other organs. We observed elevated levels of renal injury of NSG female mice bearing human TNBC tumor formed by the MDA231 cell line **(Fig. 8A)**. This was associated with increased collagen content in the renal tissue as assessed by picrosirius red staining, indicating increased levels of interstitial fibrosis. Western blot analysis confirmed these findings, showing elevated levels of fibrotic and EMT markers, including tenascin C, TGF-β, alpha-smooth muscle actin (α-SMA), β-catenin, vimentin, and snail **(Fig. 8B)**. To rule out the specificity of these pathological alterations to the human MDA231 cell line in NSG mice, we evaluated the kidney tissue of BALB/c female mice bearing the mouse breast cancer cell line 4T1. We observed similar renal pathology, consistent with the findings from the human breast cancer model **(Fig. 8C, 8D)**. To further examine the existence of cancer-kidney crosstalk with other remote organ cancers, we assessed the renal tissue of B6;129 male mice bearing a subcutaneous tumor formed by the mouse melanoma (B16) tumor. Despite the development of large tumors, there were no signs of injury or fibrosis in the renal tissue **(Fig. 8E, S2)**. Together, these findings support the existence of cancer-kidney crosstalk in certain types of organ cancer, highlighting the potential for similar pathological interactions beyond lung cancer.

**Figure 8.**
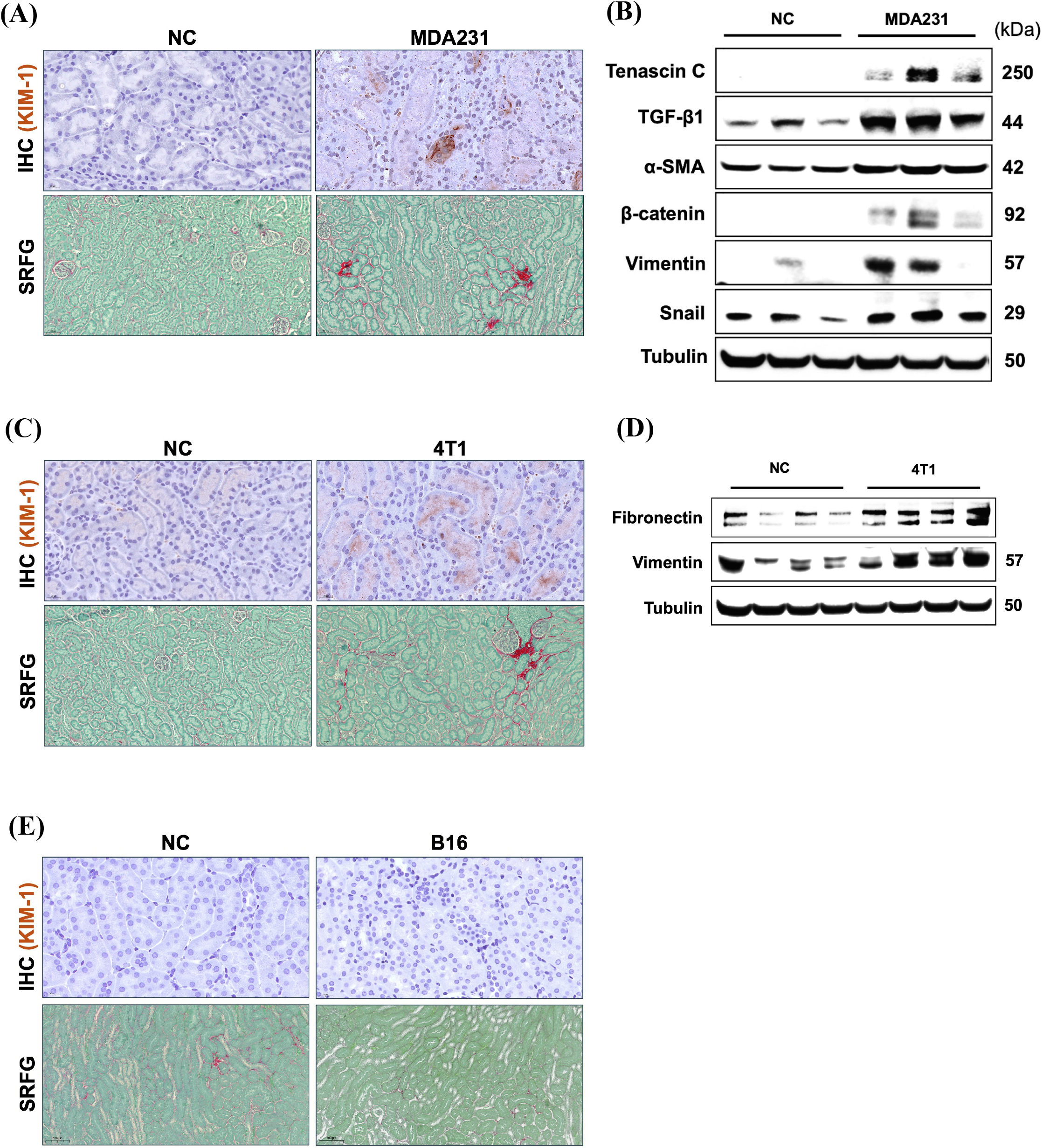
Human and mouse breast cancer induce kidney injury and fibrosis. **(A)** Representative images of IHC staining of KIM-1 and PSR staining of kidney sections of NSG female mice with or without human breast cancer (MDA231). Scale size: 50 µm and 20 µm. **(B)** Western blot analysis. **(C)** Representative images of IHC staining of KIM-1 and PSR staining of kidney sections of balb/c female mice with or without mouse breast cancer (4T1). Scale size: 50 µm and 20 µm. **(D)** Western blot analysis. **(E)** Representative images of IHC staining of KIM-1 and PSR staining of kidney sections of b6,129 male mice with or without mouse melanoma (B16).

## Discussion

In recent years, the increasing prevalence of oncology patients experiencing concurrent renal complications has drawn increased attention to the field of onconephrology. Epidemiological studies estimate that one-year incidence of AKI in oncology patients ranges from 11% to 20% [35, 36]. AKI not only contributes to significant morbidity, but also increases the risk of progression to CKD. It has been reported that 12% to 25% of patients with solid organ malignancies develop CKD [4, 37]. The onset of either AKI or CKD in this patient population constitutes a critical clinical milestone that exacerbates both morbidity and mortality independently of the cancer’s stage. This considerable incidence highlights the nephrotoxic nature of certain antineoplastic therapies, yet it prompts critical questions about whether the reported kidney complication is solely attributable to cancer therapy or if the underlying malignancy directly or indirectly impacts the kidney, contributing to nephrotoxicity. Our previous study demonstrated that metastatic Kras^G12D^ Trp53^KO^ NSCLC not only exacerbates the severity of cisplatin nephrotoxicity but also drives kidney injury, inflammation, and fibrosis in the absence of cisplatin treatment, indicating lung cancer-kidney crosstalk [19]. In the current study, we determined whether the phenomenon observed in our previous study is specific to the cancer cell line and type employed. Therefore, we assessed kidney function and physiology in mice subcutaneously injected with distinct cell lines representing various metastatic potential of Kras^G12D^ Trp53^KO^ NSCLC [32]. The highly metastatic cell line, 238N1, was associated with elevated levels of kidney injury, fibrosis, and impaired renal function compared to the less metastatic cell lines with the same driver mutations. This suggests that the severity of cancer-induced nephrotoxicity is dependent on the metastatic nature of Kras^G12D^ Trp53^KO^ NSCLC. We previously demonstrated a direct positive correlation between the tumor size of this metastatic NSCLC cell line 238N1 and the degree of kidney damage and fibrosis before and following cisplatin therapy [19]. These observations are in line with a case-control study involving patients with primary lung cancer, where the advanced tumor stage was significantly associated with increased incidence of kidney damage and higher mortality rates [38]. Notably, more than 30% of NSCLC patients are diagnosed at locally advanced stages due to the challenges in early-stage diagnosis. Therefore, there is a critical need to identify biomarkers and interventions to detect early risk of renal complications and treat or prevent progressive deterioration in kidney function in this population.

Given the increasing evidence of the bidirectional causality between acute lung injury and acute kidney injury, we aimed to investigate whether the crosstalk between the NSCLC and the kidney was due to metastasis to the lung, which could be causing kidney injury subsequent to lung injury [30, 39, 40]. Both the highly metastatic cell line 238N1 and the low metastatic cell line 394T4 were intravenously administered via the tail vein to seed the cancer cells directly into the lungs. Both the high and low metastatic lines caused substantial and similar tumor formation in the lungs and induced significant levels of kidney injury and inflammation compared to mice subcutaneously injected with these same cancer cells. However, only mice with the highly metastatic NSCLC 238N1 cell line induced kidney fibrosis. Notably, the level of kidney injury was significantly higher in mice intravenously injected with metastatic lung cancer compared to those subcutaneously injected with these same cancer cell lines. However, there was no significant difference in the level of interstitial fibrosis between the two groups. These observations suggest that the highly aggressive 238N1 NSCLC is sufficient to induce kidney fibrosis independent of lung injury. Our findings align with an observational study reporting that chronic kidney damage is significantly more frequent in patients with lung cancer compared to those with other pulmonary disorders [38].

To further examine the existence of cancer-kidney crosstalk and the specificity of this phenomenon, we evaluated kidney function and physiology in the presence of various types of remote organ cancers with distinct driver mutations in different mouse strains. Our findings suggest that tumors derived from human KRAS^G12C^ and EGFR-mutant lung cancer induce pathological crosstalk with the kidney in female and male mice of two different strains. Additionally, human and mouse breast cancer tumors were found to cause significant nephrotoxicity in female mice of two different strains. In contrast, there were no significant pathological alterations in the kidney tissue of mice with LLC or melanoma, regardless of the tumor size. This is inconsistent with a previous study that showed LLC and melanoma cause moderate renal dysfunction [41]. This could be attributed to differences in the mouse strains used or the number of cells subcutaneously injected into the animals (2×10^6^ LLC and B16 cancer cells in C57BL/6 mice compared to our 1×10^6^ LLC in B6,129 mice). This discrepancy highlights the necessity of optimizing and analyzing in detail each cancer model. It also emphasizes the need to investigate the underlying mechanisms driving the observed susceptibility to nephrotoxicity and fibrosis associated with certain cancer types across different mouse strains.

Our study suggests the existence of cancer-kidney crosstalk in a cancer type-specific manner. Importantly, our findings emphasize the critical need for incorporating different cancer types and stages in preclinical models of antineoplastic-induced nephrotoxicity. This approach would help enhance the potential for translational success in developing nephroprotective agents. Additionally, our study highlights the significance of establishing common standards to facilitate collaboration between oncologists and nephrologists at the onset of cancer diagnosis. The ability to identify susceptible oncology populations and estimate the risk of specific renal complications may significantly impact the effectiveness of cancer patients’ management.

## Acknowledgements

The authors would like to thank all of the staff at the Department of Lab Animal Research at the University of Louisville for their essential work in the care of the research animals utilized in the enclosed studies.

## Figure Legends

**Figure S1. (A)** Experimental design of lung cancer model in B6129 mice subcutaneously, injected with 1 × 10^6^ of Lewis lung carcinoma cells (LLC). **(B)** Images of tumors following euthanasia. **(C)** Tumor volume and **(D)** weight. **(E)** Level of BUN and **(F)** NGAL. **(G)** mRNA level of Kim-1, **(H)** Tnf-α, **(I)** Ccxl1, **(J)** Mcp-1, **(K)** Timp-1 and **(L)** Col1a1.

**Figure S2. (A)** Experimental design of lung cancer model in B6129 mice subcutaneously, injected with 1 × 10^6^ of melanoma cancer cells (B16). **(B)** Images of tumors following euthanasia. **(C)** Tumor volume and **(D)** weight. **(E)** Level of BUN and **(F)** NGAL. **(G)** mRNA level of Kim-1, **(H)** Tnf-α, **(I)** Ccxl1, **(J)** Mcp-1, **(K)** Timp-1 and **(L)** Col1a1.

## References

1. Rosner, M.H., et al., Onconephrology: The intersections between the kidney and cancer. CA Cancer J Clin, 2021. 71(1): p. 47–77.

2. Gupta, S., P. Gudsoorkar, and K.D. Jhaveri, Acute Kidney Injury in Critically Ill Patients with Cancer. Clin J Am Soc Nephrol, 2022. 17(9): p. 1385–1398.

3. Gudsoorkar, P., et al., Acute Kidney Injury in Patients With Cancer: A Review of Onconephrology. Adv Chronic Kidney Dis, 2021. 28(5): p. 394-401.e1.

4. Ciorcan, M., et al., Chronic kidney disease in cancer patients, the analysis of a large oncology database from Eastern Europe. PLoS One, 2022. 17(6): p. e0265930.

5. Soares, M., et al., Prognosis of critically ill patients with cancer and acute renal dysfunction. J Clin Oncol, 2006. 24(24): p. 4003–10.

6. Bermejo, S., et al., Immunotherapy and the Spectrum of Kidney Disease: Should We Individualize the Treatment? Front Med (Lausanne), 2022. 9: p. 906565.

7. Fofi, C. and F. Festuccia, Onconephrology: A New Challenge for the Nephrologist. Contrib Nephrol, 2021. 199: p. 91–105.

8. Meraz-Munoz, A., et al., Acute Kidney Injury in the Patient with Cancer. Diagnostics (Basel), 2021. 11(4).

9. Finkel, K.W. and J.R. Foringer, Renal disease in patients with cancer. Nat Clin Pract Nephrol, 2007. 3(12): p. 669–78.

10. Lam, A.Q. and B.D. Humphreys, Onco-nephrology: AKI in the cancer patient. Clin J Am Soc Nephrol, 2012. 7(10): p. 1692–700.

11. Launay-Vacher, V., et al., Prevalence of Renal Insufficiency in cancer patients and implications for anticancer drug management: the renal insufficiency and anticancer medications (IRMA) study. Cancer, 2007. 110(6): p. 1376–84.

12. Yonezawa, A., [Platinum agent-induced nephrotoxicity via organic cation transport system]. Yakugaku Zasshi, 2012. 132(11): p. 1281–5.

13. Madias, N.E. and J.T. Harrington, Platinum nephrotoxicity. Am J Med, 1978. 65(2): p. 307–14.

14. Coca, S.G., S. Singanamala, and C.R. Parikh, Chronic kidney disease after acute kidney injury: a systematic review and meta-analysis. Kidney Int, 2012. 81(5): p. 442–8.

15. Chawla, L.S., et al., The severity of acute kidney injury predicts progression to chronic kidney disease. Kidney Int, 2011. 79(12): p. 1361–9.

16. Chawla, L.S., et al., Acute kidney injury and chronic kidney disease as interconnected syndromes. N Engl J Med, 2014. 371(1): p. 58–66.

17. Amdur, R.L., et al., Outcomes following diagnosis of acute renal failure in U.S. veterans: focus on acute tubular necrosis. Kidney Int, 2009. 76(10): p. 1089–97.

18. Kovesdy, C.P., Epidemiology of chronic kidney disease: an update 2022. Kidney Int Suppl (2011), 2022. 12(1): p. 7–11.

19. Orwick, A., et al., Lung cancer-kidney cross talk induces kidney injury, interstitial fibrosis, and enhances cisplatin-induced nephrotoxicity. Am J Physiol Renal Physiol, 2023. 324(3): p. F287–f300.

20. Oliver, T.G., et al., Chronic cisplatin treatment promotes enhanced damage repair and tumor progression in a mouse model of lung cancer. Genes Dev, 2010. 24(8): p. 837–52.

21. Ranasinghe, R., M.L. Mathai, and A. Zulli, Cisplatin for cancer therapy and overcoming chemoresistance. Heliyon, 2022. 8(9): p. e10608.

22. A genomics-based classification of human lung tumors. Sci Transl Med, 2013. 5(209): p. 209ra153.

23. Dogan, S., et al., Molecular epidemiology of EGFR and KRAS mutations in 3,026 lung adenocarcinomas: higher susceptibility of women to smoking-related KRAS-mutant cancers. Clin Cancer Res, 2012. 18(22): p. 6169–77.

24. El Osta, B., et al., Characteristics and Outcomes of Patients With Metastatic KRAS-Mutant Lung Adenocarcinomas: The Lung Cancer Mutation Consortium Experience. J Thorac Oncol, 2019. 14(5): p. 876–889.

25. Winslow, M.M., et al., Suppression of lung adenocarcinoma progression by Nkx2-1. Nature, 2011. 473(7345): p. 101–4.

26. Sharp, C.N., et al., Subclinical kidney injury induced by repeated cisplatin administration results in progressive chronic kidney disease. Am J Physiol Renal Physiol, 2018. 315(1): p. F161–F172.

27. Sharp, C.N., et al., Repeated administration of low-dose cisplatin in mice induces fibrosis. Am J Physiol Renal Physiol, 2016. 310(6): p. F560–8.

28. Yuan, Q., et al., A Klotho-derived peptide protects against kidney fibrosis by targeting TGF-β signaling. Nat Commun, 2022. 13(1): p. 438.

29. Alge, J., et al., Two to Tango: Kidney-Lung Interaction in Acute Kidney Injury and Acute Respiratory Distress Syndrome. Front Pediatr, 2021. 9: p. 744110.

30. Husain-Syed, F., A.S. Slutsky, and C. Ronco, Lung-Kidney Cross-Talk in the Critically Ill Patient. Am J Respir Crit Care Med, 2016. 194(4): p. 402–14.

31. Thies, K.A., et al., Pathological Analysis of Lung Metastasis Following Lateral Tail-Vein Injection of Tumor Cells. J Vis Exp, 2020(159).

32. Rashid, O.M., et al., Is tail vein injection a relevant breast cancer lung metastasis model? J Thorac Dis, 2013. 5(4): p. 385–92.

33. Fragiadaki, M. and R.M. Mason, Epithelial-mesenchymal transition in renal fibrosis - evidence for and against. Int J Exp Pathol, 2011. 92(3): p. 143–50.

34. Sheng, L. and S. Zhuang, New Insights Into the Role and Mechanism of Partial Epithelial-Mesenchymal Transition in Kidney Fibrosis. Front Physiol, 2020. 11: p. 569322.

35. Salahudeen, A.K., et al., Incidence rate, clinical correlates, and outcomes of AKI in patients admitted to a comprehensive cancer center. Clin J Am Soc Nephrol, 2013. 8(3): p. 347–54.

36. Christiansen, C.F., et al., Incidence of acute kidney injury in cancer patients: a Danish population-based cohort study. Eur J Intern Med, 2011. 22(4): p. 399–406.

37. Kitchlu, A., et al., Perspectives From an Onconephrology Interest Group: Conference Report. Can J Kidney Health Dis, 2020. 7: p. 2054358120962589.

38. Pedersen, L.M. and N. Milman, Prevalence and prognostic significance of proteinuria in patients with lung cancer. Acta Oncol, 1996. 35(6): p. 691–5.

39. Li, X., F. Yuan, and L. Zhou, Organ Crosstalk in Acute Kidney Injury: Evidence and Mechanisms. J Clin Med, 2022. 11(22).

40. Herrlich, A., Interorgan crosstalk mechanisms in disease: the case of acute kidney injury-induced remote lung injury. FEBS Lett, 2022. 596(5): p. 620–637.

41. Xu, W., et al., A novel antidiuretic hormone governs tumour-induced renal dysfunction. Nature, 2023. 624(7991): p. 425–432.

