## Supplementary Figures for "Remote organ cancer adversely alters renal function and induces kidney injury, inflammation, and fibrosis"

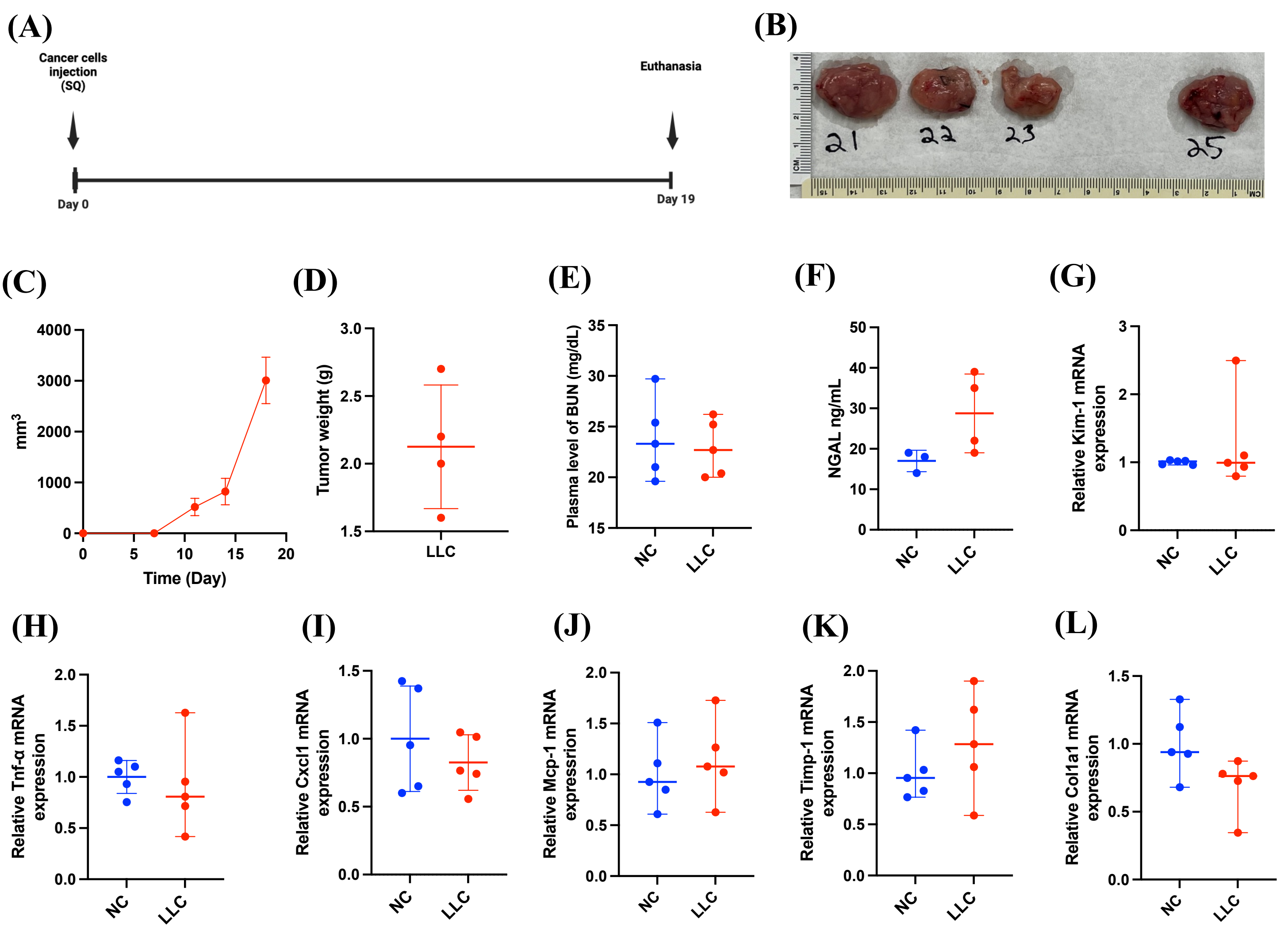


**Figure S1. (A)** Experimental design of lung cancer model in B6129 mice subcutaneously, injected with 1 × $10^6$ of Lewis lung carcinoma cells (LLC)**. (B)** Images of tumors following euthanasia. **(C)** Tumor volume and **(D)** weight**. (E)** Level of BUN and **(F)** NGAL. **(G)** mRNA level of Kim-1, **(H)** Tnf-α**, (I)** Ccxl1, **(J)** Mcp-1**, (K)** Timp-1 and **(L)** Col1a1**.**

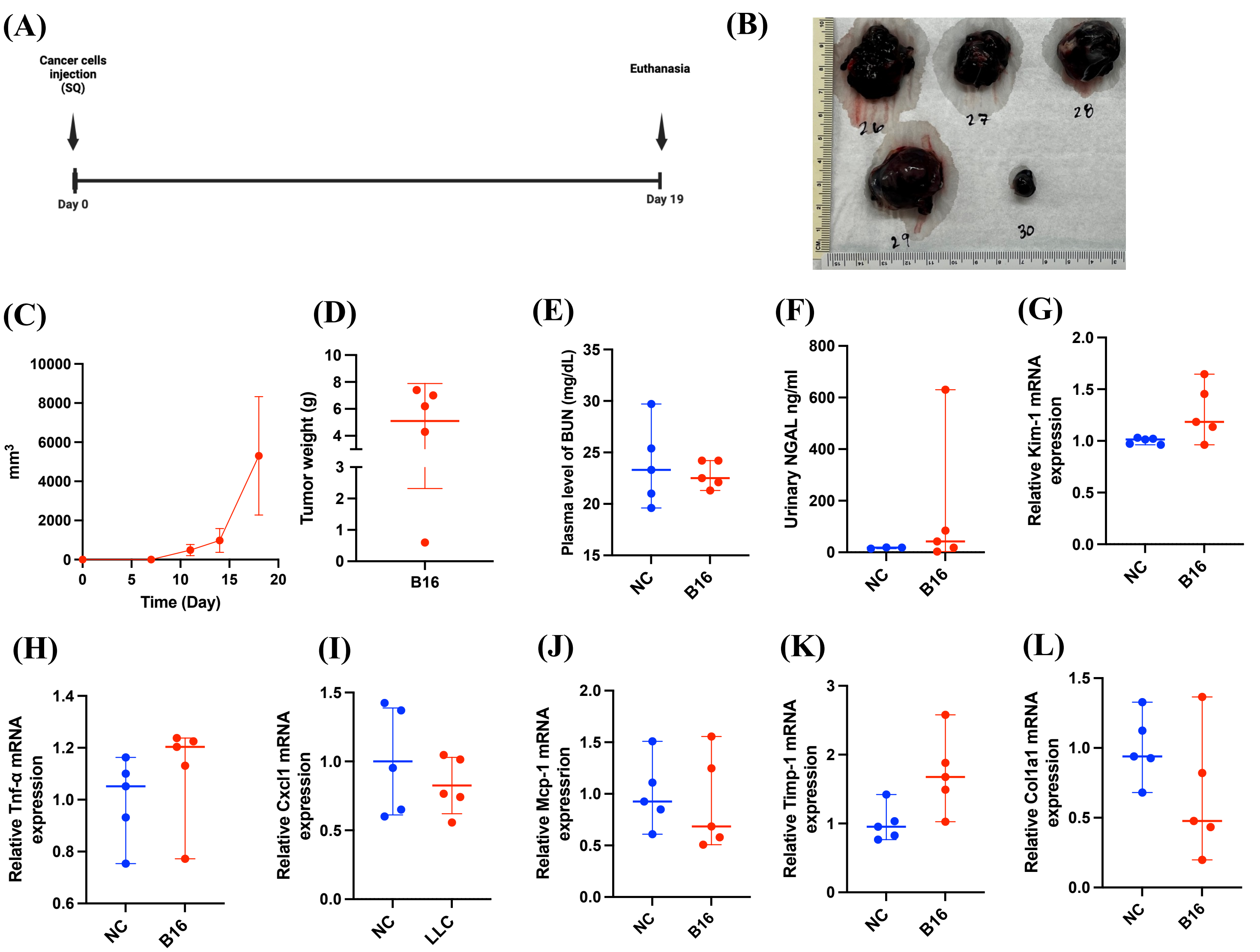


**Figure S2. (A)** Experimental design of lung cancer model in B6129 mice subcutaneously, injected with 1 × $10^6$ of melanoma cancer cells (B16). **(B)** Images of tumors following euthanasia. **(C)** Tumor volume and **(D)** weight. **(E)** Level of BUN and **(F)** NGAL. **(G)** mRNA level of Kim-1, **(H)** Tnf-α**, (I)** Ccxl1, **(J)** Mcp-1**, (K)** Timp-1 and **(L)** Col1a1**.**
